# From migrants to residents: Genomic insights into adaptive strategies in European robins (*Erithacus rubecula*)

**DOI:** 10.64898/2026.06.26.734870

**Authors:** Corinna Langebrake, Georg Langebrake, Javier Pérez-Tris, Juan Carlos Illera, Miriam Liedvogel

## Abstract

Bird migration evolved as an adaptation to seasonally changing habitats. Migratory behaviour can vary within the same species in case of partial migratory behaviour, i.e. one population (or individual) is migratory and another one is resident. Species that exhibit a wide variety of migratory phenotypes provide valuable systems to understand the evolutionary drivers behind different phenotypes and how populations adapt to habitats with distinct seasonality. The European robin (*Erithacus rubecula*) expresses migratory behaviour in central and northern areas of the species distribution range, whereas populations in the South and on the Macaronesian islands are predominantly resident, providing a suitable system to investigate these questions. We use high coverage whole genome re-sequencing data of 125 European robins to investigate how migration behaviour affects population structure and demography, and how it affects the selection landscape in the genome. Genetic structure in European robins coincides with migratory phenotype and geography and populations are characterised by distinct demographic histories. Our results suggest that both the continental resident population as well as the Macaronesian island populations have derived independently from an ancestral migratory population. Unexpectedly, tests for differential selection revealed extensive positive selection pressure acting across all chromosomes in the resident populations, while selective sweeps are largely absent from migrants. We speculate that this might be an analytical artifact due to mismatching timescales between what population genomics methods can detect and the scale on which migration behaviour likely evolved in the robin. We suggest that future studies on the genomics of migration should more focally account for different time scales on which these processes happen, such as including the wider phylogenomic background of the target species, to capture the full evolutionary history of migratory traits.

## Introduction

The evolution of avian migration on a macro scale (i.e. millions of years ago) is hypothesised to be mainly driven by seasonality of the habitat and breeding site fidelity (Salewski & Bruderer, 2007; Winger *et al*., 2019). The dispersal theory combines the Northern-and Southern-home theory (migration evolved either out of the tropics by dispersal or from Northern habitats through higher seasonality; Levey & Stiles, 1992; Rappole *et al*., 1992; P. Bell, 2000) by stating that migratory behaviour could have evolved both close and far from the equator (Salewski & Bruderer, 2007). The theory suggests that migration evolves by either stepwise dispersal into more seasonal habitats (i.e. in terms productivity such as food availability) or a climatic change in the breeding habitat, leading to unsuitable conditions during a predictable time of the year. Birds evolved different strategies to cope with seasonal environments, of which migration behaviour is just one (Winger *et al*., 2019). Others include a change in foraging strategy and food storage, growing denser or thinner feathers, physiological adaptations such as dropping body temperature during the night, group roosting and actual hibernation in the common poorwill (*Phalaenoptilus nuttalli*; Jaeger, 1948, 1949; Hohtola, 2012; Blix, 2016; Osváth *et al*., 2018).

Seasonal migration is a behaviour that combines the orchestration of many different traits, resulting in an accurate response to intrinsic and extrinsic conditions (Piersma *et al*., 2005; Liedvogel *et al*., 2011). These traits would include orientation, timing, propensity, hyperphagia and nocturnal activity (Zugunruhe). We know that these traits are heritable and thus have a genetic basis in many species (Helbig, 1991; Berthold *et al*., 1992; Pulido & Berthold, 1998). The manifestation of the migratory phenotype is very species-specific, and in some cases even population specific (Delmore *et al*., 2020a, 2020b; Sokolovskis *et al*., 2023) or varying within one breeding population (Adriaensen & Dhondt, 1990; Hegemann *et al*., 2015; Zúñiga *et al*., 2017; Acker *et al*., 2021).

On the population scale, it is hypothesised that the expression of a migratory phenotype is described by the threshold model (Pulido, 2011): Migratory propensity or liability is considered to be a normally distributed trait underlying the binary expressed phenotype (resident/migrant). A threshold determines when the migratory phenotype is expressed and the individual’s liability and its relative position to the threshold determines if a migratory or resident phenotype is exhibited. For facultative partial migrants (i.e., migratory phenotype is flexible within one individual) it is assumed that individuals with a liability close to the threshold can change their migratory strategy based on intrinsic and/or extrinsic conditions. The threshold model implies that if a population is close to the threshold, all individuals have the genomic machinery to express both phenotypes. If a population is exclusively migratory, all individuals are too far from the threshold to rapidly switch to the alternate phenotype, even with drastically changing environmental conditions. If this specialised migratory state persists, over (a yet undetermined) timespan and with increasing distance to the threshold, phenotypic plasticity decreases through genetic assimilation/canalisation and switching back would require evolutionary change (Pulido, 2011). The subset of genes evolved/optimised for the adaptations to a migratory phenotype should be positively selected for in migrants or under purifying selection to maintain optimal variants. However, selection pressure on genomic features that specialised for migratory behaviour should be relaxed in residents to a certain extent. If a population is exclusively resident and positioned far from the threshold, we would expect genetic canalisation to increase and in consequence phenotypic flexibility to decrease. Thus, at some point, populations are no longer able to rapidly change their phenotype as a (plastic) reaction to changing environmental conditions and in extreme cases of e.g. flight loss the adaptations to residency are irreversible.

While migratory behaviour in some (predominantly long-distance migratory) species, such as the Northern wheatear (*Oenanthe oenanthe*), hardly varies between different populations across the species distribution range (i.e. strong canalisation; Bairlein *et al*., 2012; Winger *et al*., 2019), other species show extremely high phenotypic variability on a population level. In these taxa, phenotypic variability allows for rapid adaptation to changing conditions. An iconic example is the Eurasian blackcap (*Sylvia atricapilla*), which exhibits the full range of phenotypic variation from long-distance migration to resident populations (Delmore et al., 2020a); and in addition to variation in the propensity and distance migrated, also exhibits variation in migratory direction (Delmore *et al*., 2020a, 2020b, 2023). To further understand mechanisms of how migratory behaviour or residency evolves on a population level we need additional species showing similar responses to add a comparative perspective.

To investigate the loss or gain of migration behaviour on a population level as a consequence of local seasonality differences, we focus on multiple migratory and resident populations of the European robin (*Erithacus rubecula*), with populations on the Iberian Peninsula showing migration and residency in geographic proximity. Additionally we include a migratory population in Germany as well as resident populations on the Macaronesian islands. Such a scenario is well suited to characterise genomic signatures of the focal behaviour as differentiation due to geographic distance can be controlled for, while potential effects of canalisation should be evident, which makes it an ideal system to study the effects of migration loss/gain on population structure, demography and selection pressure. Also, due to a similar setting as in Delmore *et al*. (2020a), our results are directly comparable with the blackcap system. We use high-coverage whole genome re-sequencing (WGS) data to investigate how robins became adapted to seasonal habitats, and characterise how selection pressure acts on phenotypically distinct populations.

The ancestral migratory phenotype has not been conclusively demonstrated for continental European robins. However, it is plausible to assume that given their close relationship to sedentary African forest robins (Zhao *et al*., 2023) and two newly recognised *Erithacus* species on the Canary Islands (*E. marionae* and *E. superbus*) that diverged early from the most recent common ancestor (MRCA, ca 3 and 2 mya, respectively), the ancestral continental robin might have been resident (Valente *et al*., 2017; Fjeldså *et al*., 2020; Sangster *et al*., 2022). Thus, it remains an open question whether the continental resident robins in the Iberian Peninsula emerged through migration drop-off from northern populations or whether continental robins gained the migratory phenotype after expanding to more seasonal habitats. We aim to distinguish between these scenarios by investigating demographic differences between the populations.

## Results

### Clear separation between migrants and residents

We sequenced 125 individual robins (84 males and 41 females) sampled in 17 different locations across Europe (Figure 1a) and called variants to a reference genome (GenBank GCA_903797595.1, individual information in Table S1), resulting in 27,501,388 single nucleotide variants after filtering. One individual was removed due to high missingness and five additional individuals were excluded from further analysis due to relatedness.

**Figure 1.**
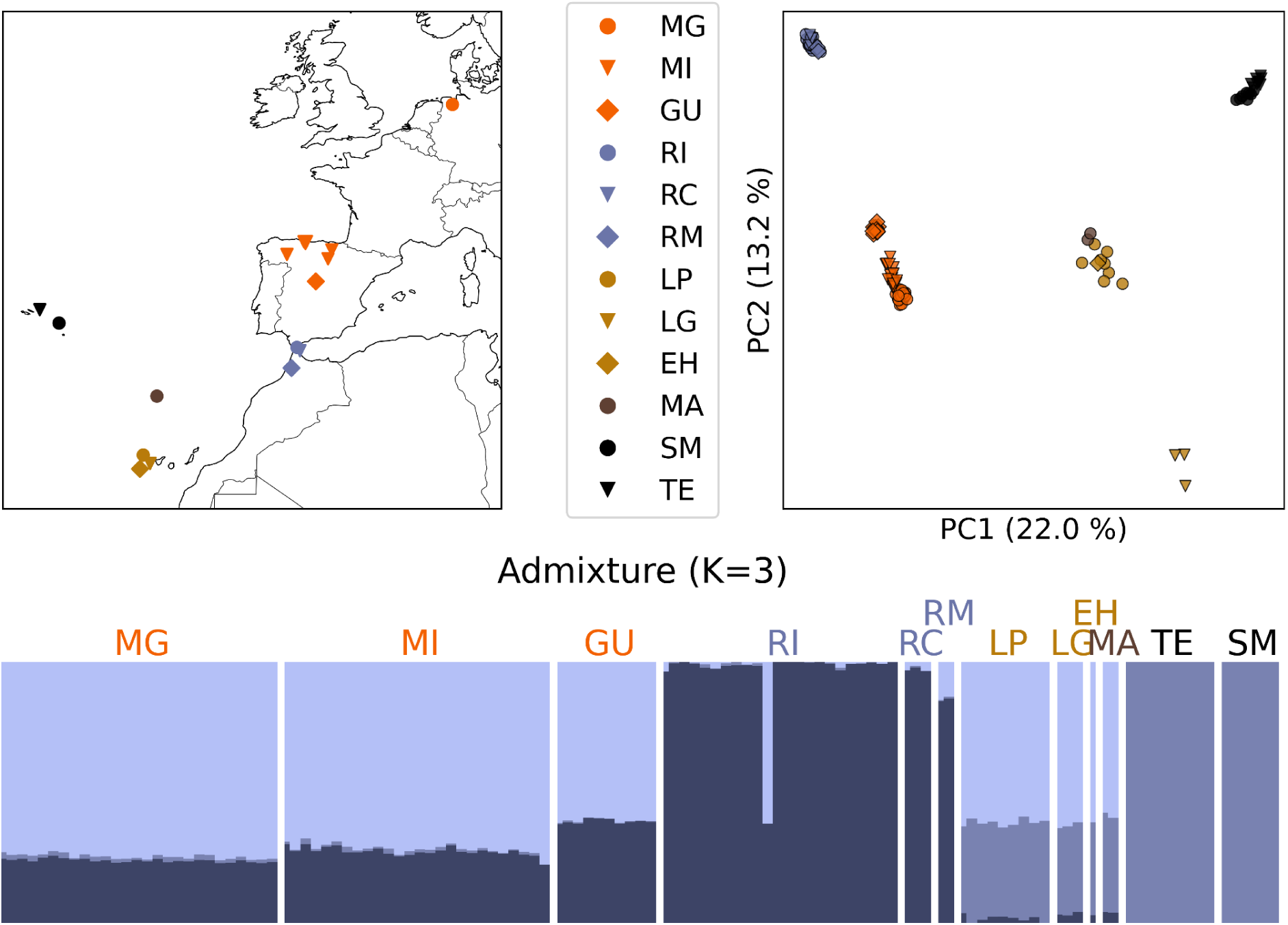
Population structure of resident and migratory European robins. **a)** Individuals were sampled from migratory populations (orange) in Germany (light, MG), northern Spain (middle, MI) and central Spain (dark, GU), as well as resident populations on the European continent (blue) in southern Spain (light, RI), Ceuta (middle, RC) and Morocco (dark, RM). Additional resident populations were sampled on the Macaronesian Islands (brown) from the Canaries (light, CA), Madeira (middle, MA) and the Azores (dark, AZ) b) Samples cluster according to their geographic Origin in the PCA, with one axis separating continental residents from the migrants and another separating the macaronesian islands from the migrants. c) Using 3 genetic clusters best explains the genetic structure in an admixture analysis. The full names of locations are explained in the Methods and Table S1.

Population clustering, admixture analysis and a phylogeny identify clear population separation between all continental migrants, continental residents, the Canary Islands together with Madeira and the Azores. Admixture analysis suggests an optimum of 3 clusters, with one cluster corresponding to the continental residents, which is partially shared by the migrants, one to the Azores and partially the Canaries and Madeira and one cluster connecting Canaries and Madeira with the migrants (Figure 1c). The PCA further corroborates an independent split of both resident groups from the modern migrants with Macaronesian residents mostly separating on one axis from the migrants, while the continental residents separate on a perpendicular axis (Figure 1b). The phylogenetic tree again follows the same pattern, suggesting that ancestral diversity is maintained in the migrant population, while the resident populations have independently derived from that ancestral population (Figure 2). Bootstrap support for most branches is low, as within populations there is little genetic structure, but branches with high support match the picture painted by the other methods. Not only are the resident populations clearly delineated, additional structure within the populations is also made clear. On the Macaronesian islands all methods separate the Azores from Canaries and Madeira, with the latter being genetically more similar to the migratory population. Additionally, La Gomera is separated from the rest of the Canaries and Sao Miguel and Terceira separate both in the PCA and the phylogeny. In the continental residents, clear separation can be seen between Africa and Europe in the phylogeny, though this is not evident from the PCA and Admixture. Additionally, individuals from Guadarrama, the migratory population that is geographically closest to the southern residents, shows genetic differentiation from the rest of the migratory population and acts as a distinct sister taxon to the southern residents. The individuals from the North Coast that were phenotyped as resident/uncertain are not separated from other Northern migrant individuals and cluster by geography and not migratory behavior. Given the phenotype characterisation we cannot exclude these individuals being partial migrants. One individual from the southern sample site clusters in all of the analyses within the GU individuals (northern migrants). This specific individual was sampled in mid-August, which is post-breeding, but should still be before migration onset (Remisiewicz, 2002). Thus, this could be evidence of genetic exchange between these populations, however, there is no admixture in any of the other southern individuals, which makes an unexpected early dispersal or migration event of this particular individual more likely.

**Figure 2.**
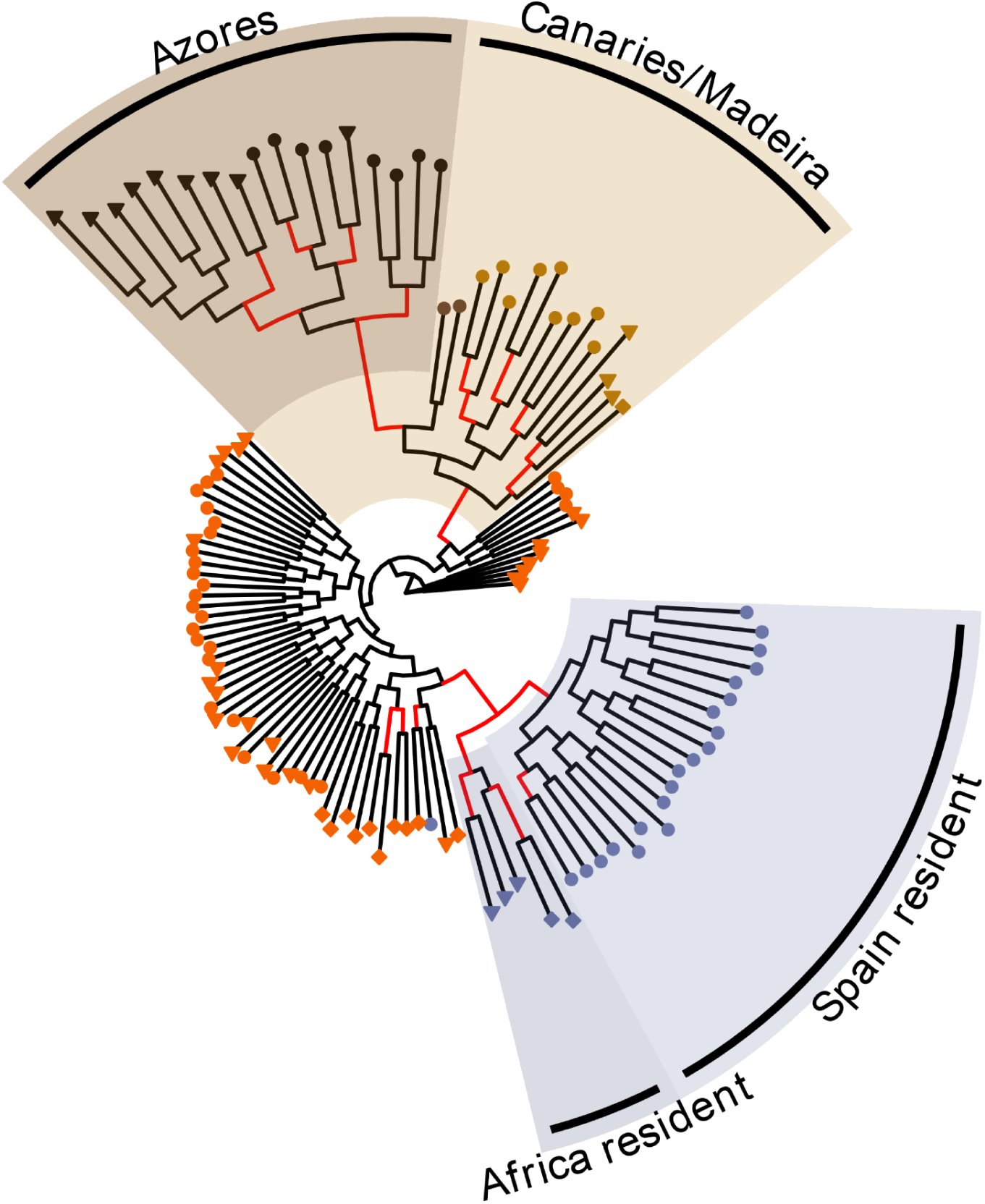
Maximum Likelihood phylogeny of European robins. Tree of all individuals used in this study generated using maximum likelihood. Leaf colors indicate sampling population (light orange - Germany; orange - Northern Spain; dark orange - central Spain; light blue - southern Spain; blue - Ceuta; dark blue - Morocco; light brown - Canaries; brown - Madeira; dark brown - Azores). Branches present in at least 95% of bootstrap iterations are highlighted in red.

### Demography and inbreeding

The inferred demographic history shows a pattern typical for temperate avian populations (Nadachowska-Brzyska *et al*., 2015), with a population expansion before, and a population reduction during the last glacial period (LGP, 115,000 to 11,700 kya; Figure 3). Cross-coalescence rate analysis shows that continental residents split off from the last common ancestor around the start of the LGP and show a stronger population reduction during the LGP compared with the rest of the robins. Just after the LGP, about 100 kya later, the Macaronesian island populations split off with an immediate strong decrease in population size. The modern migrant population however expands after the LGP, matching its vastly larger range compared to the resident populations examined in this study. Note however that recent estimates are generally less reliable.

**Figure 3.**
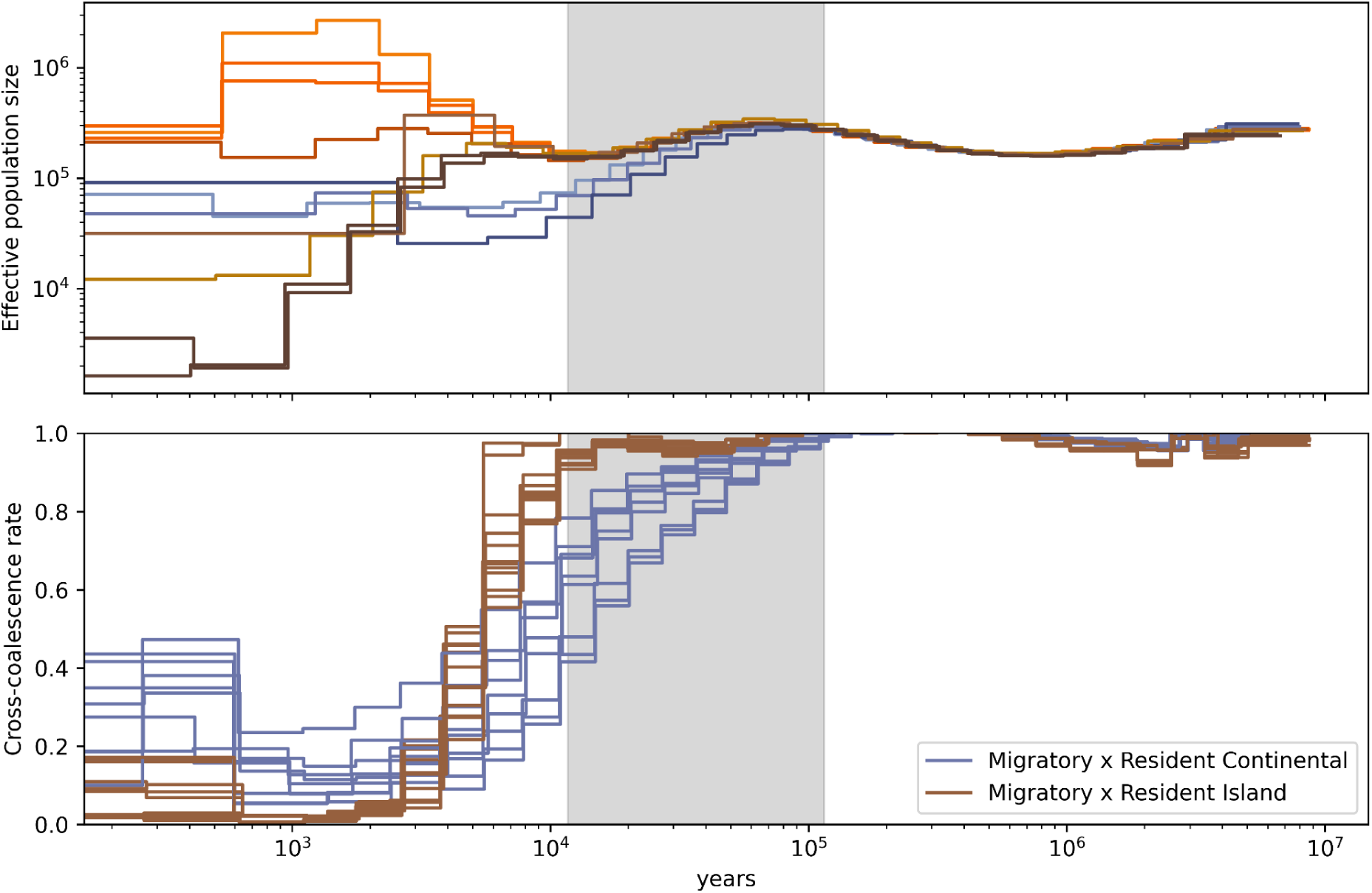
Demographic history of European robins. a) Effective population size (N_e_) was inferred from 5 individuals per population randomly picked across all sample locations light orange - Germany, orange - Northern Spain, dark orange - central Spain, light blue - southern Spain, blue - Ceuta, dark blue - Morocco, light brown - Canaries, brown - Madeira, dark brown - Azores). Grey shading indicates the last glacial period (LGP) from 115.000 to 11.700 years ago. b) Cross coalescence rate (CRR) was calculated for every pair of populations in our dataset. Depicted are the CCRs between migrant and resident populations. A CCR of 1 indicates no separation between populations while a CCR of 0 indicates completely split populations. Continental residents and migrants started to split at the beginning of the LGP while Macaronesian residents only split from the migrant populations at the end of the last LGP.

Homozygosity in all resident groups is significantly higher compared to migrant groups and higher than would be expected under Hardy-Weinberg equilibrium, in line with a significantly higher degree of inbreeding in these populations (T-test, all p < 0.01; Figure S1). For migrants, the mean observed (OH) and mean expected heterozygosity (EH) are not different.

Since analyses revealed no apparent genetic differentiation within geographically close sampling locations, further analysis focus on four groups based on genetic and geographic clusters: birds from the Wold form the German migrants (GM), northern Iberian locations were combined as Iberian migrants (IM), all continental resident populations were combined as continental residents (CR) and La Gomera, La Palma and Madeira were combined as island residents (IR). Because of limitations in sample sizes from the Azores and El Hierro, these populations were excluded from further analysis.

### Population differentiation

We characterised population differentiation between migrants and residents through weighted F_ST_ and Delta F_ST_ in 50 kb windows. Delta F_ST_ accounts for linked selection and local adaptation by controlling for F_ST_ in non-target (migrant-migrant and resident-resident) comparisons (Vijay *et al*., 2016). We identified a number of regions that are classified as outliers in multiple population comparisons, including 4 windows that are identified in all 4 focal comparisons. When looking at differentiation that is specific to one population (i.e. differences found comparing a single resident population to both migratory populations), it becomes clear that most identified regions are specific to the resident populations (see figure S2). With 184 windows identified in at least 2 focal comparisons we decided to only further discuss regions found to be under selection with different methods.

### Positive selection in southern residents is in contrast to no signs of selection in northern migrants

As the populations only recently started and continue to diverge, this allows us to detect more recent selection events related to this event. We used xp-EHH (cross-population Extended Haplotype Homozygosity) statistics (Sabeti *et al*., 2007; Gautier *et al*., 2017) for differential selection analysis in both resident and migrant populations. xp-EHH identifies hard selective sweeps based on haplotype homozygosity and cross-population comparisons control for local variation in recombination rate while genome-wide normalization of between-population haplotype length controls for different population histories (Sabeti *et al*., 2007). A stark contrast of selection pressure was identified in the two phenotypes with hardly any sweeps in migrants and several sweeps in residents, in line with the results of the delta F_ST_ analysis (Figure 4, 5, S3). Furthermore, there was a strong difference in the number of windows under selection in CR compared to IR (Figure S3). In addition to detecting selection based on haplotype homozygosity, we also used hapFLK (Fariello *et al*., 2013) to identify divergent haplotype blocks in two or more populations, indicative of recent selective sweeps. This approach is complementary to the above analysis as selection is detected based on population differentiation (F_ST_-like). It controls for both genetic drift within populations by controlling for N_e_ as well as hierarchical structuring of populations. Combined, delta F_ST_, xp-EHH and hapFLK agree on 2 regions under differential selection between continental residents and both migratory groups and 4 regions between residents on the Canaries and Madeira and both migratory groups (most likely driven by positive selection as negative selection acts homogenizing; Table 1). The strongest signal was found in continental residents on chr29 in a long haplotype block (ca. 200 kb), containing 14 genes. This region is also characterised by low recombination and Tajima’s D and high Delta F_ST_ (Figure 6). Similar to what was shown in Eurasian blackcaps, all identified outlier regions were under selection in resident populations (Delmore *et al*., 2020a).

**Figure 4:**
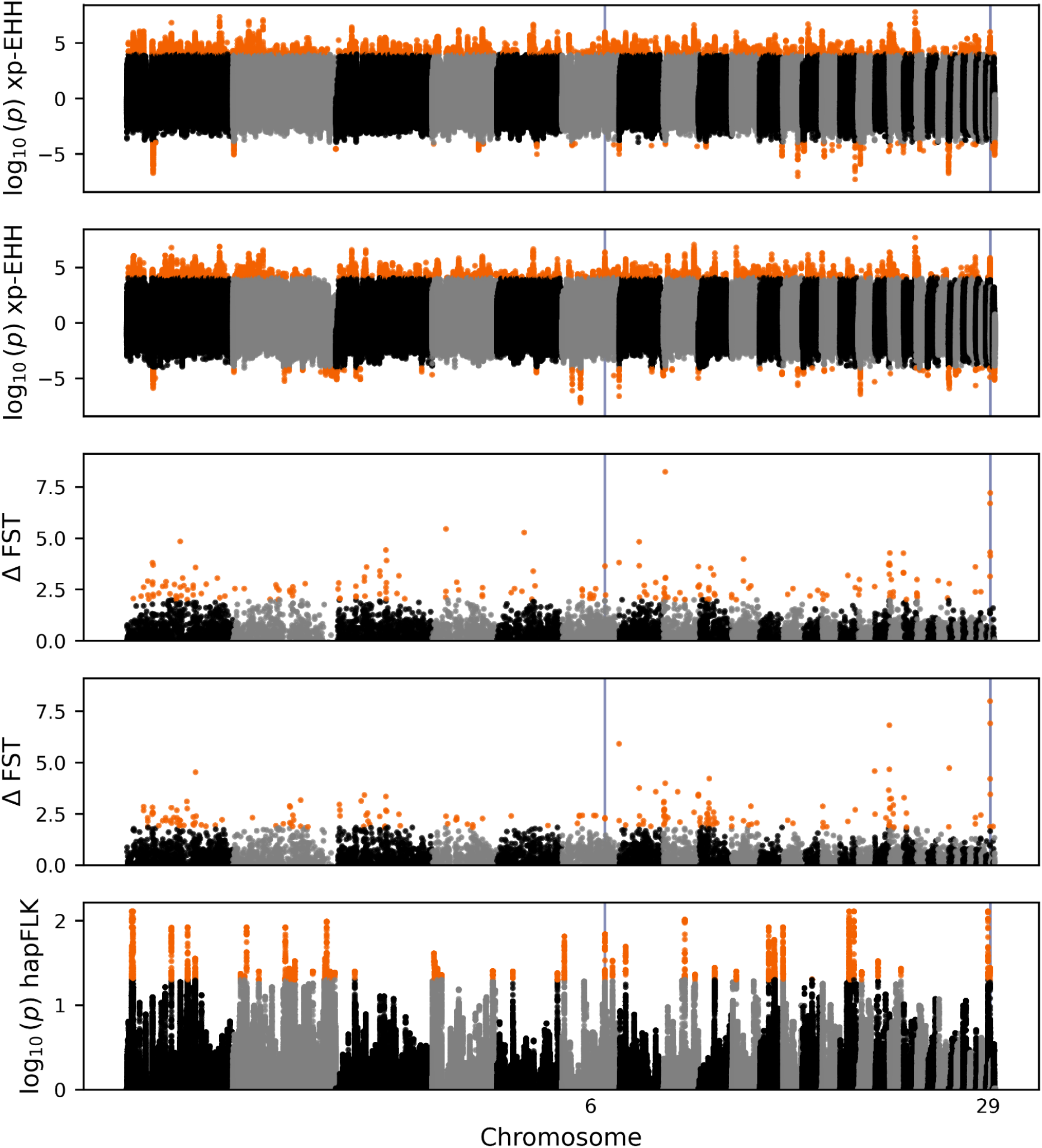
Population differentiation and selection signatures between migratory and continental resident robins. a) Divergent selection between continental residents (CR) and Iberian migrants (IM) identified by xp-EHH. Positive values indicate selection in CR, negative values indicate selection in IM. Highlighted values in orange show values for which the corrected p value is below 0.05. b) xp-EHH values between CR and German migrants (GM). Positive values indicate selection in CR, negative values selection in IM. c) z-transformed Delta-F_ST_ of the comparison CR-IM corrected for CR-island residents and IM-GM. Orange outliers correspond to windows in the upper 99th percentile. d) z-transformed Delta-F_ST_ of the comparison CR-GM corrected for both CR-island residents and IM-GM. e) Divergent selection test based on genetic differentiation (hapFLK, haplotype extension of F_ST_-based FLK). Outlier values shown in orange are identified based on an FDR corrected p-value of less than 0.05.

**Figure 5:**
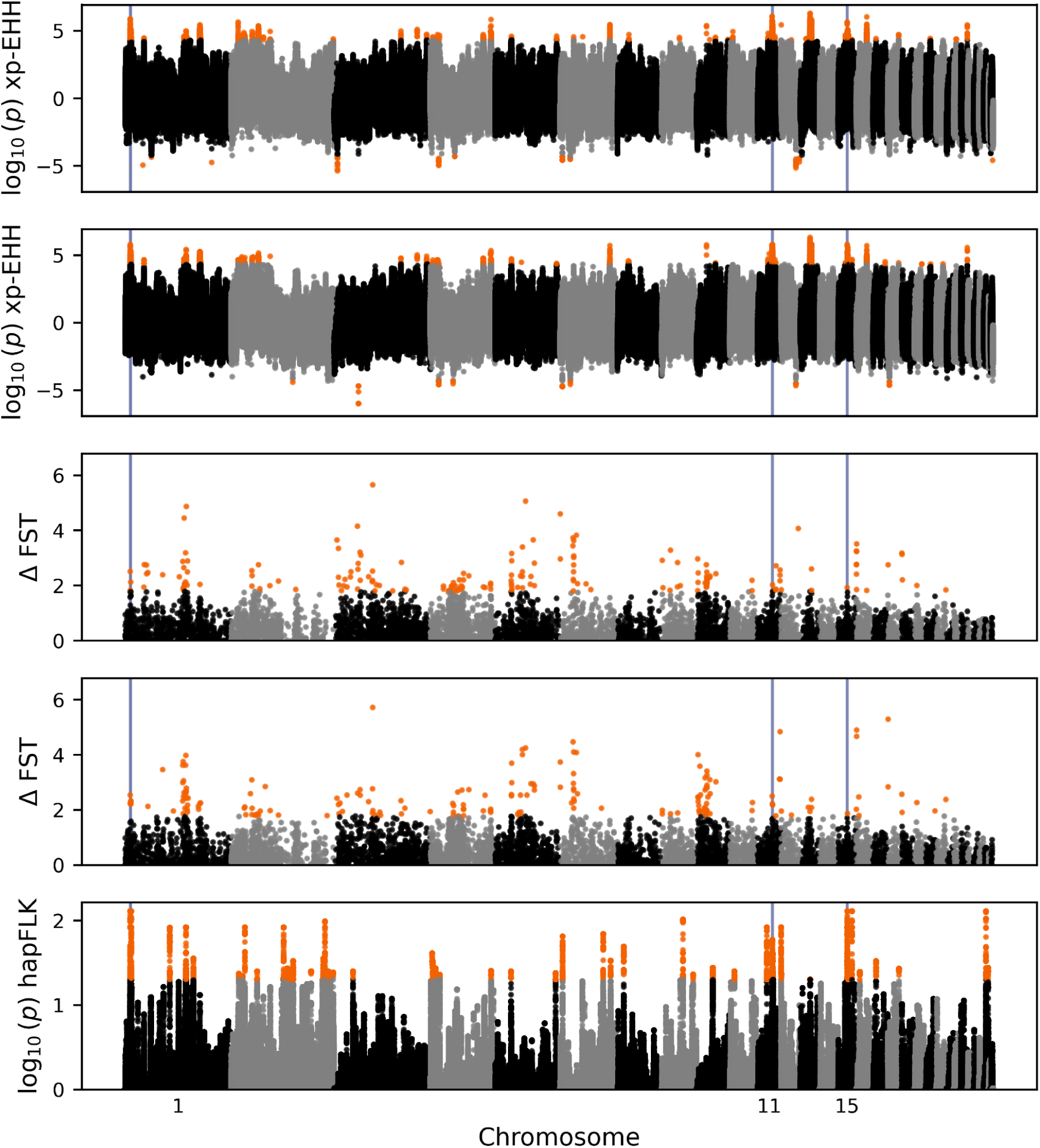
Population differentiation and selection signatures between migratory and island resident robins. a) Divergent selection between island residents (IR) and Iberian migrants (IM) identified by xp-EHH. Positive values indicate selection in IR, negative values indicate selection in IM. Highlighted values in orange show values for which the corrected p value is below 0.05. b) xp-EHH values between IR and German migrants (GM). Positive values indicate selection in IR, negative values selection in IM. c) z-transformed Delta-F_ST_ of the comparison IR-IM corrected for IR-continental residents and IM-GM. Orange outliers correspond to windows in the upper 99th percentile. d) z-transformed Delta-F_ST_ of the comparison IR-GM corrected for both IR-continental residents and IM-GM. e) Divergent selection test based on genetic differentiation (hapFLK, haplotype extension of F_ST_-based FLK). Outlier values shown in orange are identified based on an FDR corrected p-value of less than 0.05.

**Figure 6.**
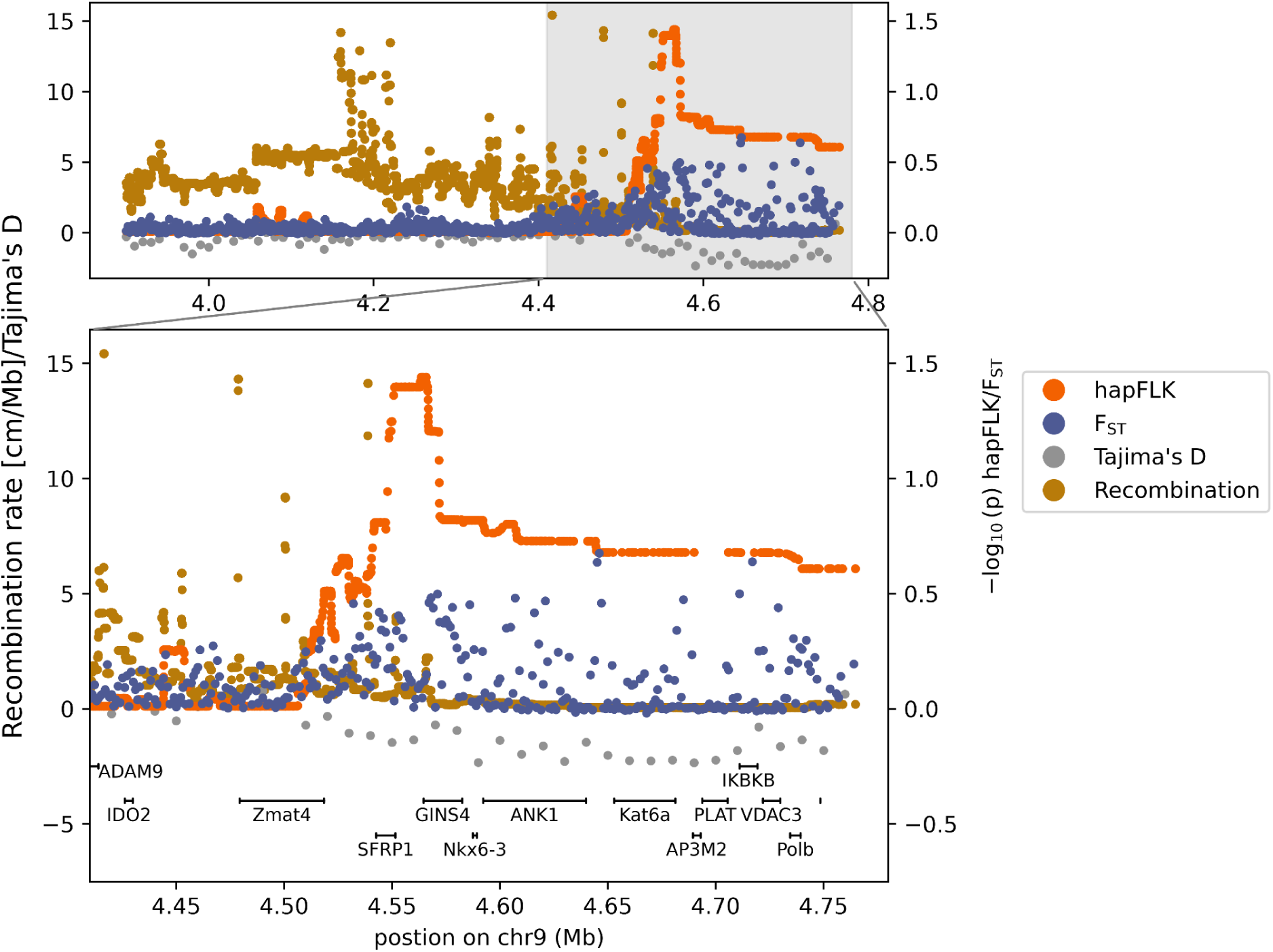
Candidate region on chr29. A haplotype block of ca 200 kb on chr29 is characterised by high hapFLK p-values (black) and elevated weighted FST in 10 kb non-overlapping windows for N-S comparisons (pink). Recombination rate (rec, cM/Mb, blue) and Tajima’s D (purple) are reduced. 14 genes are located in this differentiated region (the region highlighted in light grey is magnified in the lower section).

**Table 1.**
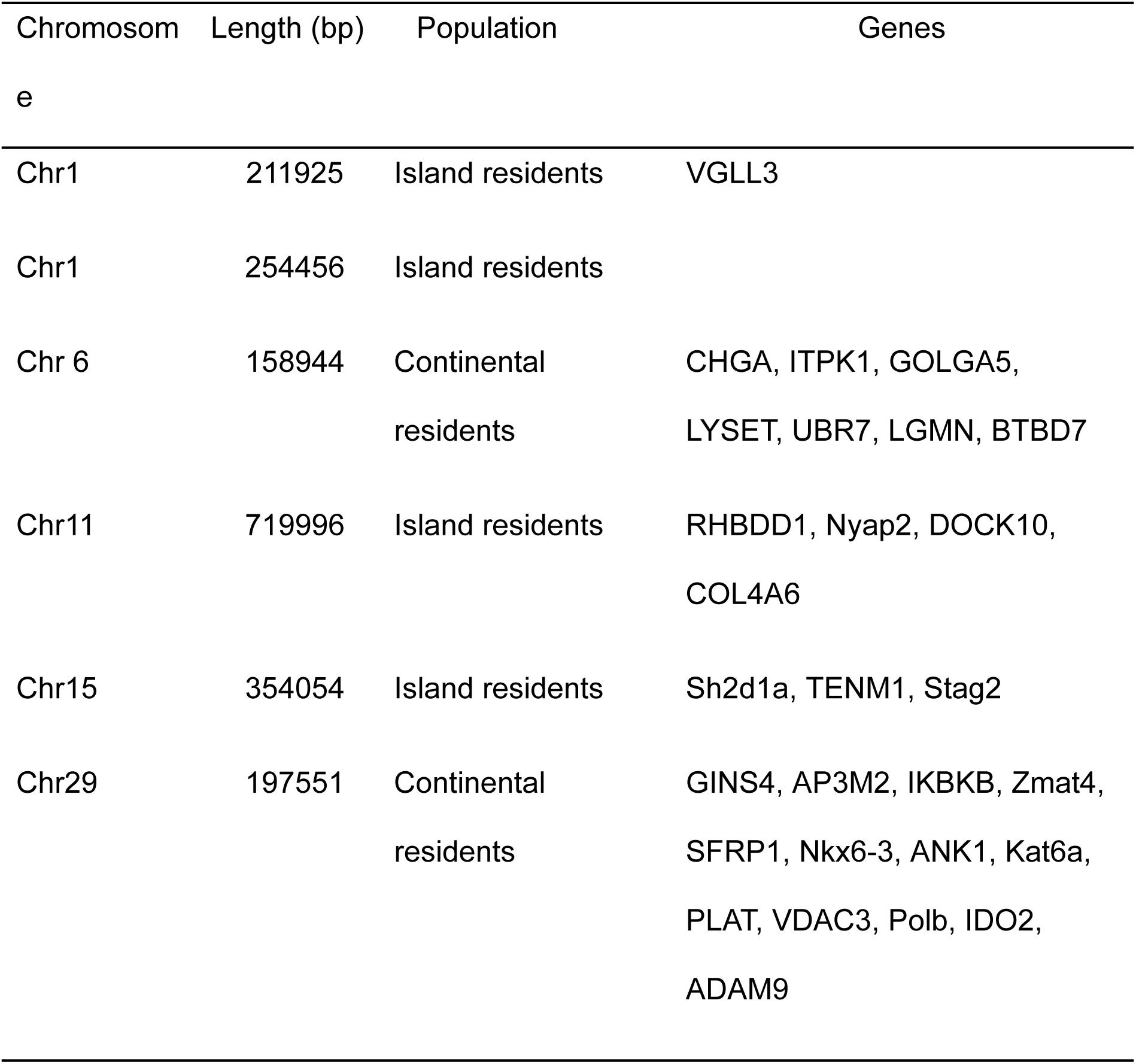
Identified candidate regions under divergent selection (hapFLK). Resident southern robins were compared with migratory northern robins using hapFLK. We report chromosome location and the length of the selected region as well as list the population in which the selection signal was found. Annotated genes within 50 kb of the identified sweep are listed.

In the chr29 peak, genes of various functions identified in mouse or human model systems with potential relevance to migratory behaviour can be found, including a major negative regulator of the WnT pathway (SFRP1, Secreted Frizzled Related Protein 1) which regulates obesity (Lagathu *et al*., 2010), and a transcription factor involved in development of the central nervous system (NKX6-3, NK6 homeobox 3). AP3M2 (Adaptor Related Protein Complex 3 Subunit Mu 2), another protein in the peak is involved in protein trafficking to lysosomes and has been associated with abnormal accumulation of lipids in cells and tissue in humans, which in migratory birds is particularly interesting in the context of hyperphagia as preparation for successful migration. Further, VDAC3 (Voltage-dependent anion-selective channel protein 3), a voltage-dependent anion channel is located in this region, which in knock-out experiments in mice results in non-functional spatial learning (Weeber *et al*., 2002). Mice with knocked-out VDAC3 were not able to learn optimal routes through a maze and behaved like naive mice of the untrained control group. Multiple genes in the peak were also related to eye development and indicated in diseases causing blurry vision (IDO2, Zmat4) and night blindness (ADAM9). Interestingly, two genes found in selective sweeps in continental residents - CHGA (chromogranin A, Chr6) and PLAT (tissue plasminogen activator, Chr29) - are functionally linked as part of the same autocrine feedback system regulating catecholamine release at neuroendocrine cells (Parmer *et al*., 2000). The physiological relevance of catecholamine regulation during migration is underscored by Ivy *et al*. (2025), who demonstrated that adrenaline increases markedly during high-altitude migratory flight in yellow-rumped warblers (*Setophaga coronata*), with the magnitude of the response reflecting the degree of hypoxic stress experienced.

Most genes found in regions under selection in the island residents are involved in neuronal development (NYAP2, Neuronal Tyrosine-Phosphorylated Phosphoinositide-3-Kinase Adaptor 2), postsynapse assembly (DOCK10, Dedicator Of Cytokinesis 10) and neural connectivity (TENM1, Teneurin Transmembrane Protein 1). VGLL3 has been shown to be critical for the repression of adipocyte enhancers in mice, and in humans a correlation with blood cholesterol levels, insulin resistance, and body mass index was identified (Seol *et al*., 2026).

## Discussion

### The ancestral European robin was likely migratory

Our population structure analyses conclusively show that after the LGP, around 10 kya ago, the contemporary migratory population was able to spread throughout all of Europe, likely as a consequence of post glacial range expansions and re-colonisation of previously unsuitable habitat (Hewitt, 2000), whereas the continental resident population has kept a stable population size after the LGP. It seems plausible that the migratory phenotype facilitated the opportunity to expand into habitats with suitable breeding conditions that were not accessible to obligate residents due to cold winters. This is also observed in contemporary populations where migratory robins are expanding their breeding ranges north (Günther *et al*., 2026).

During the LGP however, both robin populations present show a population decline. This fits to what has been previously reported for a variety of temperate residents or short distance migrants (Nadachowska-Brzyska *et al*., 2015), but is in contrast to three more recent studies on Eurasian blackcaps (Delmore *et al*., 2020a), northern pied flycatchers (*Ficedula hypoleuca*; Nadachowska-Brzyska *et al*., 2016) and red-backed shrikes (*Lanius collurio*; Thorup *et al*., 2021), representing migratory species with the potential to cross the Sahara. These species seem to maintain growing populations over the last glaciation period until present, possibly reflecting the ability to track suitable habitat as a consequence of the migratory phenotype. The European robin is a short distance migrant, likely not capable of crossing the Sahara Desert, which had less precipitation during the last glaciation compared to today (Collins *et al*., 2017), and thus acts as a full ecological barrier (Illera *et al*., 2008). There is evidence that migratory behaviour was maintained during the LGP, especially in trans-Saharan migrants (Ponti *et al*., 2020; Somveille *et al*., 2020). Maintaining migration during the LGP could have benefited the modern robin migratory population, potentially explaining the clear difference we find in population reduction between the migratory and the resident population during that time. Other factors that could influence different trends between populations and species could be driven by glacial refugia location and habitat specialisation, also with respect to declining populations in the more specialised collared and Atlas flycatcher during the LGP (Nadachowska-Brzyska *et al*., 2016).

Given that the ability to migrate was very likely present at the end of the LGP and a gain of migration seems highly unlikely during the LGP, while a maintenance is possible and could even be beneficial, we infer that migration was likely ancestral to the populations investigated in this study. On a deeper timescale however, extensive and comparable taxon sampling is needed to robustly characterise the ancestral phenotype of European robins. Especially in the context of two newly recognised species on the Canary Islands (split 3 and 2 mya), it is possible that the ancestral phenotype of continental migrants was indeed resident when they diverged from their closest relatives about 1 mya (Valente *et al*., 2017; Sangster *et al*., 2022) and potentially evolved a migratory phenotype as a consequence of dispersal into more seasonal habitats (Salewski & Bruderer, 2007). The possibility of a deeper resident ancestry is further plausible given that the closest relatives of European robins are the mostly resident African forest robins (Zhao *et al*., 2023).

Our demographic results also suggest that the Macaronesian islands were colonised by the modern migratory population around the end of the LGP (about 100 ky after the first migratory drop-off), coinciding with the range expansion of continental migrants. This finding is consistent with the hypothesis that migration, and its subsequent loss following island colonisation, has played a key role in shaping extant island bird assemblages (Dufour *et al*., 2024). Interestingly, the Azorean populations appear to be differentiated from those of Madeira and the Canary Islands, with the latter archipelagos emerging as the most likely source populations for the colonisation of the Azores. This pattern is unusual because the prevailing north-easterly and north-westerly trade winds are expected to facilitate avian dispersal from north to south across Macaronesia (Illera *et al*., 2012). To our knowledge, only two other bird species show a comparable south-to-north colonisation pattern: the island canary, *Serinus canaria* (Dietzen *et al*., 2006), and Berthelot’s pipit, *Anthus berthelotii* (Illera *et al*., 2007; Martin *et al*., 2023).

### Signs of genomic canalisation in two migratory dropoffs over time

The two likely migratory drop-off events we observe have very different characteristics: (i) The more recent colonisation of the Macaronesian islands (about 10 kya), most likely involving comparatively few individuals, appears to be a consequence of post glacial range expansions, while (ii) the split of continental residents around 100 kya likely reflects a different mechanism related to population separation in glacial refugia. Speciation and population differentiation as a consequence of multiple glacial cycles and survival in glacial refugia is a common phenomenon in various bird species in the Northern hemisphere (Avise & Nelson, 1989). The cross-coalescence rate between islands and migrants suggest a quick separation (see Fig. 3b), with likely no or low gene transfer after the initial colonisation. On the other hand, the older separation of continental residents from the ancestral population is much slower and population separation appears not yet complete. This is further indicated by the genome-wide genetic similarity of the central Iberian migratory population with the continental residents, suggesting some amount of gene flow.

The vast majority of selective sweeps driving both migration drop-off events occurred in the resident populations. However, the events differ in the amount of selection observed. In the older split of the continental residents there are more windows identified to be under differential selection in xp-EHH compared to the macaronesian populations (Figure S3). The different selection intensity in the two drop-off events could reflect the different divergence time scales and consequently varying intensity of genomic canalisation with respect to the threshold model of migration (Pulido, 2011). As both resident populations are obligatory resident, major behavioural changes may require only a few genomic changes. Selection observed in residents may therefore reflect not the behavioural change towards residency, but rather the elimination of costly migratory genomic features that evolved in a migratory ancestor (such as excessive fat deposition, i.e. hyperphagia, in preparation for a successful migratory journey), or the adaptation to local conditions (i.e. less seasonal and more specific habitat characteristics). As a consequence of this canalisation, rapid shifts to a migratory phenotype are likely no longer possible and require genomic changes (Pulido, 2011).

In addition to the two stages of canalisation presented here (old split vs recent split), European robins have partial migratory populations in central Europe. These would represent a state of no canalisation, where only few (i.e. those “switching off” migration) to no (exclusively environmental triggers) genetic differences between migrants and residents exist. Combined, these three systems would provide a unique opportunity to study how loss of migration evolves over time.

### No parallel adaptation to residency on various timescales

The identified peaks harbour genes that could be involved in the migratory phenotype. Extended haplotype blocks of highly differentiated and apparently linked regions similar to the peak on chr29 were also identified in other migratory systems, explaining a high percentage of migratory trait variation differentiating different subspecies of willow warblers (*Phylloscopus trochilus*; Sokolovskis *et al*., 2023), Swainson’s thrushes (*Catharus ustulatus*; Delmore *et al*., 2016) and European quails (*Coturnix coturnix*; Sanchez-Donoso *et al*., 2022). It was discussed that genes encoding complex traits are under linked selection pressure (for migration e.g. physiological and behavioural changes such as fat accumulation, nocturnal activity) and thus might be inherited as a gene package (Liedvogel & Lundberg, 2014; Delmore *et al*., 2016; Justen & Delmore, 2022), eventually resulting in supergenes, which could facilitate the rapid adaptation of complex traits. It was argued for Swainson’s thrushes that a candidate region for this gene package was identified (Delmore *et al*., 2016), however, no overlap of this region was found with other systems such as willow warbler (Sokolovskis *et al*., 2023) or European quail (Sanchez-Donoso *et al*., 2022). The region on chr29 shows some aspects of such a gene package and it provides a candidate region that should be investigated in other populations and closely related species.

None of the identified candidates matches genes identified in other studies focussing on migration genomics, though targeted pathways overlapped. However, most other studies did not focus on migration propensity but on other traits (e.g., direction) and thus the most comparable system to our study here is the Eurasian blackcap (Delmore *et al*., 2020a) and common yellowthroat (Zamudio-Beltrán *et al*., 2025). Thus the apparent problem that arises from population-level analyses of species with a distinct species specific evolutionary history is that apparently almost no common genetic basis for the trait under consideration (e.g. migratory propensity) can be identified (Swainson‘s thrush: Delmore *et al*., 2016; blue/golden-winged warbler: Toews *et al*., 2019; European blackcap: Delmore *et al*., 2020a; peregrine falcon: Gu *et al*., 2021; common quail: Sanchez-Donoso *et al*., 2022; willow warbler: Sokolovskis *et al*., 2023, common yellowthroat: Zamudio-Beltrán *et al*., 2025), which is in contrast to the initial hypotheses drawn out before high throughput sequencing enabled these studies (Liedvogel *et al*., 2011). However, we suggest that this apparent lack of shared loci under selection may not be surprising, as each species adapts to very distinct environmental pressures (Garcia-Porta *et al*., 2022). Furthermore, if a trait such as migration direction or timing varies on the population level (i.e., on very short time scales) it is hard to imagine how this trait could be phylogenetically constrained between species. Indeed, a recent study highlights that migratory propensity in several populations of the same species is only partially mediated by selection of the same genes (Zamudio-Beltrán *et al*., 2025) similar to the pattern in the two independent migratory drop-off events in our study system. Thus, we expect that each species and even each population finetunes its underlying machinery, targeting distinct points of signalling pathways that may regulate the same underlying biological trait (Martin *et al*., 2024; Zamudio-Beltrán *et al*., 2025). This is supported by the fact that specific pathways associated with energy metabolism, nervous system, immune response and circadian rhythm regulation are recurrently found to be associated with migratory traits (Delmore *et al*., 2016, 2020a; Toews *et al*., 2019; Gu *et al*., 2021; Sanchez-Donoso *et al*., 2022; Sokolovskis *et al*., 2023; Zamudio-Beltrán *et al*., 2025). With an increasing body of literature, only few genes associated with timing, energetics of flight and morphology are highlighted as recurring outliers in different migratory species (reviewed in Zamudio-Beltrán *et al*., 2025), indicating convergent or parallel evolution on the species level (Estandía *et al*., 2023; Martin *et al*., 2024).

### Mismatch of timing in methods and phenotype changes

The apparent lack of selection on the migratory phenotype is unexpected, especially given the assumption that migratory traits should be under strong (purifying/positive) selection pressure (Liedvogel *et al*., 2011). However, the same pattern was also found in blackcaps, where continental residents (from breeding areas matching those of resident robins studied here) also showed more selective sweeps compared to migrants (Delmore *et al*., 2020a). We propose that this apparent mismatch between expectation and observation might be due to a mismatch between the time scale we consider in a population genomics analysis, and the timescale on which the behaviour evolves. Methods to detect selection signals in populations can, depending on recombination rate, mutation rate and generation time, primarily identify recent sweeps (Sabeti *et al*., 2006; Tanaka *et al*., 2023). In humans, some methods can detect selection up to 250.000 years ago (Sabeti *et al*., 2006) and tests for long haplotypes like i-HS and xp-EHH detect very recent sweeps (up to 50.000 years). However this would be less in the songbirds discussed here as their generation time is shorter and if migratory adaptations evolved before that, they would be missed. It is assumed that migration has a common genetic basis (Liedvogel *et al*., 2011) and phylogenetic analysis suggests that the trait is constrained on a phylogenetic timescale (Dufour *et al*., 2020; Langebrake *et al*., 2024). This means that the basic machinery for successful migratory behaviour might at least partially evolve on a much slower timescale compared to what we can capture on a population genomics level (i.e. conserved on a species level). Unless two different migratory phenotypes are contrasted within the same species, such as direction, timing, winter longitude/latitude and distance, the signal would be missed. In the resident robins, we can identify selection likely associated with the new environment as well as the new, resident phenotype. If continental robins were ancestrally resident prior to ∼1 mya (Fjeldså *et al*., 2020), by expanding to the species level it would thus be more likely to identify the genomic architecture of migratory propensity. Ideally, the two newly described robin species on the Canary Islands, as well as the closest related forest robins should be included in future genomic approaches. The power of considering also the evolutionary history of the focal species was demonstrated by (Zhan *et al*., 2014): By including closely related species in the monarch butterflies system, they were able to assign a migratory ancestral phenotype to extant resident populations (Zhan *et al*., 2014). For focally studying the genomic architecture of migration we thus propose to expand the time window under consideration, as we currently do not know how “labile” the genomic basis of this behaviour really is and on which time scale it evolves (e.g. independent origin in each genus, family or order, and different migratory traits evolving on different timescales).

## Conclusion

We identify clear differences between migratory and resident robin breeding populations across continental and island populations, despite their recent divergence and potentially incomplete lineage sorting. Our results suggest two migratory drop-off events on the continent and on the Macaronesian islands, originating from a migratory ancestor and now expressing obligate resident behaviour. This behavioural adaptation to less seasonal habitats is potentially governed by only a few major loci and is reflected by selective sweeps occurring exclusively in the resident population. The absence of selection signals in the migrants might hint towards analytical constraints, with respect to the timescale on which the available tools allow us to capture adaptations necessary for the migratory phenotype. As continental European robins may have a resident ancestry, with two resident species older than the continental migrants (3 and 2 mya compared to 1 mya respectively), the inclusion of these species in subsequent studies could shed light on the specific adaptations necessary for migration behaviour in the European robin. The identified haplotype blocks could function as regulatory switches for the migration behaviour, as has been discussed for Swainson’s thrushes (Delmore *et al*., 2016). Although migration behaviour seems to be governed by extensively linked haplotype blocks in different species, these blocks are not shared between species. However, instead of rejecting the hypothesis of a homology of migration, we suggest that insights from the population level inform us on how each focal species adapts to the very specific requirements of their respective ecological niches. Thus, they modify and maintain the genomic machinery of the migratory phenotype, which does not exclude a homology of this fundamental machinery in a broader phylogenomic context. In order to identify or reject the hypothesis of a deeper homology, we need to extend our analyses to broader time scales, testing for a common machinery and signals of parallel and convergent evolution across different taxonomic levels.

## Methods

### Samples and resequencing data

We obtained blood samples from 125 European robins (*Erithacus rubecula,* 84 males and 41 females), sampled at seventeen locations across Europe from populations that were classified as either migratory or resident (for details see Table S1) based on population-level long-term observations as described in previous studies (Tellería *et al*., 2001; Pérez-Tris & Tellería, 2002; Hera *et al*., 2014; Demongin, 2016). Diagnostic methods used for classifications were ringing and capture-recapture analyses combined with morphometrics and stable isotope analysis. Genomic DNA was extracted via a standard salt extraction protocol. Libraries were prepared through the Illumina DNA PCR-free prep, tagmentation and sequenced on a NovaSeq with paired-end 150 bp reads. Each sample was sequenced on to an average coverage of 30 X (Table S1, data is available under the accession number PRJEB71790). Variant calling, including adapter trimming, mapping to the reference genome, SNP calling and filtering was performed using variant-calling v.1.0.0 (Langebrake & Weissensteiner, 2025). This snakemake (Mölder *et al*., 2025) pipeline uses fastp (Chen, 2025) for adapter trimming, bwa-mem2 (Vasimuddin *et al*., 2019) and samtools (Danecek *et al*., 2021) for read mapping and bcftools (Danecek *et al*., 2021) for variant calling and filtering. The resulting vcf file was filtered for biallelic sites. For non population structure analysis, one individual with a high proportion of missing genotypes (PN1886, northern N-population) was excluded. Filter for minor allele frequency MAF and more stringent missingness filter per genotype were applied according to the analyses carried out, see for detailed information in following sections.

### Genome annotation

Functional gene annotation of the European robin reference genome (Dunn *et al*., 2021) was performed using Maker version 3.1.4 (Campbell *et al*., 2014). This included soft masking the genome for transposable elements using RepeatMasker 4.1.2 (Smit *et al*., 2013) with a library of transposable elements from the collared flycatcher (Suh *et al*., 2018) and the blue-capped cordon bleu (Boman *et al*., 2019). For transcription level evidence we used ISOseq data from the retina, brain, muscle, lung, liver, heart and skin from a single European robin (Xu *et al*., 2025), for which circular consensus sequences were generated to yield full-length transcripts. Additionally, the transcriptome of chicken (*Gallus gallus*) was included as transcriptional evidence and all proteins in swissprot from birds were used as protein evidence. Using blast and exonerate through maker, the evidence were mapped to the reference and used as input for the ab-initio gene predictors Augustus and snap (Korf, 2004; Stanke & Morgenstern, 2005). Snap was additionally trained for two iterations, first using gene positions derived from evidence mappings and secondly using the output of the previous generation. The resulting genes were filtered for an annotation edit distance of less than 0.5, a metric provided by Maker to assess the goodness-of-fit of the annotation to the transcriptional evidence (Campbell *et al*., 2014).

### Population structure

We restricted our analyses to autosomes and calculated Tajima’s D in 1000 bp-windows and heterozygosity with vcftools (Danecek *et al*., 2011). We identified closely related individuals (parent-child or siblings) through the KING-robust kinship estimator (Manichaikul *et al*., 2010) in plink2 (Chang *et al*., 2015) and excluded five related individuals from subsequent analyses. Based on this adjusted dataset, we ran pca in plink2 and ADMIXTURE (Price *et al*., 2006; Alexander *et al*., 2009) to characterise and visualise population structure. For the latter, we explored the fit of K = 2 to 8 (K equals number of populations/clusters).

### Phasing and recombination rate

As some analyses require phased data, we performed statistically inferred phasing using phasing-snakemake v0.1.0 (Langebrake, 2026a). This snakemake pipeline uses shapeit5 (Hofmeister *et al*., 2023). This requires a recombination map which was generated using recombination-snakemake v0.1.0 (Langebrake, 2026b), a pipeline implementing fine-scale recombination rate estimation using pyrho (Spence & Song, 2019), a demography-sensitive program that infers the per-generation, per-base recombination rate “r”. This pipeline follows the procedure described in (Bascón-Cardozo *et al*., 2024). We used a single individual in msmc2-snakemake v0.1.0 (Langebrake, 2026c) to infer historic population size for input in pyrho. In this mode, no phasing is required. The mutation rate was set to 4.6 ∗ 10*^−^*9 mutation per site per generation (Smeds *et al*., 2016). The genetic map that is generated by the pipeline was used as input for phasing-snakemake.

### Demography

We used msmc2-snakemake v0.1.0 (Langebrake, 2026c). This pipeline uses MSMC2 (Schiffels & Wang, 2020) for demography inference on a population level based on phased SNPs. First, we treated each sample location as a “population” and included not more than 5 individuals per population, with random subsampling in case of more individuals. We corrected for the mutation/recombination ratio by setting-r to 1.5. This was calculated by taking the average recombination rate of the whole genome as inferred through pyrho with an assumed mutation rate of 4.6∗10*^−^*9 mutation per site per generation (inferred from data on flycatcher, (Smeds *et al*., 2016)). To calculate the generation and population size, a mutation rate of 4.6 ∗ 10*^−^*9 and a generation time of two years were used.

### Differentiation

We used the PBS-snakemake pipeline v0.1.0 (Langebrake, 2026d) to calculate delta F_ST_ in 50 kb non-overlapping windows. The pipeline calculates F_ST_ using vcftools between every pair of populations specified and uses this output to calculate delta F_ST_ for focal comparisons by first normalising F_ST_, followed by removing the maximum of normalised F_ST_ in non-focal comparisons for every window. This controls for local adaptation (Vijay *et al*., 2016). Focal comparisons were all pairwise F_ST_ between the groups GM, IM and CR, IR, for non-focal comparisons we used the F_ST_ between GM - IM and CR - IR.

### Selection

We used two different approaches to infer differential selection pressure in the northern migrants and southern residents. The method xp-EHH (cross-population Extended Haplotype Homozygosity) statistics (Gautier *et al*., 2017) can detect recent selective sweeps based on long-range haplotype homozygosity as a measure of linkage disequilibrium in each of two populations. The cross-population comparison controls for variation in recombination and genome-wide normalization of between-population haplotype length controls for different population histories (Sabeti *et al*., 2007). We used the snakemake pipeline rehh-snakemake v0.1.0 (Langebrake, 2026e) which uses the R-package rehh (Gautier *et al*., 2017) to calculate xp-EHH for every focal comparison between migrants and residents. The resulting test statistic is directional, so values were either kept as is or multiplied by-1 so that values > 0 mean selection in the residents and values < 0 correspond to selection in the migrants. P-values are FDR corrected in the pipeline and sites with a corrected p-value < 0.05 were defined as outliers.

We also used a different approach to detect differential selection based on the hapFLK method (Fariello *et al*., 2013). This algorithm accounts for hierarchical structure of populations and genetic drift within populations by controlling for N_e_ and is the haplotype extension of the F_ST_-based (genetic differentiation) FLK statistic. For this we used the snakemake pipeline hapflkl-snakemake v0.1.0 (Langebrake, 2026f). The pipeline can automatically find the optimal number of clusters using fastPHASE and a cross-validation procedure, which in our case turned out to be at 25. We tested for differential selection between all four focal groups (GM, IM, CR, IR). The population covariance matrix was calculated based on the whole genome and was used for hapFLK calculation in the next step. P-values are false-discovery-rate corrected as part of the pipeline and we identified sites as under differential selection with a corrected p-value < 0.05.

We focussed on peaks where all 3 methods (delta FST, xp-EHH and hapFLK) detected an outlier region. Outlier regions were defined as belonging to the same peak if they were within 50kb of an outlier region in a different method. With two focal comparisons for every group we selected those regions for every population that were present in both focal comparisons involving the given population. This was also used together with xp-EHH to infer directionality of selection. We used our annotation to pinpoint genes within the selective sweeps identified with hapFLK and panther (Thomas *et al*., 2022) to identify the associated GO terms.

## Acknowledgements

We thank Arne Nolte and Raphael Schween for providing IsoSeq data which were used for genome annotation. We thank the Max Planck Society (MPRG grant MFFALIMN0001 to ML) and the DFG (DFG, German Research Foundation) for funding of projects Z02 and Nav05 within SFB 1372 – Magnetoreception and Navigation in Vertebrates (project number 395940726 to ML) and under Germany’s Excellence Strategy - EXC 3051/1 „NaviSense” (project number 533653176 to ML). JCI was funded by two research grants from the Spanish Ministry of Science, Innovation and Universities, and the European Regional Development Fund (PGC2018-097575-B-I00; PID2022-140091NB-I00). JPT was funded by grant PID2020-116121GB-I00 funded by MCIN/AEI/10.13039/501100011033 and by “ERDF A way of making Europe”. Computational resources were provided by the ROSA Cluster at the Carl-von-Ossietzky University, Oldenburg, supported by the DFG and the Ministry for Science and Culture of Lower Saxony.

## Permits

The Regional Government of Andalusia (SGYB/AF/FJRH; Re-35-36/13), Asturias (2013/001891), Diputación Foral de Álava (Exp 08/32), Madrid (10/160876.9/10), and Castilla y León (EP/CYL/33/2013), the National Parks of Sierra Nevada (ENSN/BRL/MCF, Expte.Inv.: 3/12) and Picos de Europa (CO/09/012/2013) gave permission to conduct fieldwork and blood sampling. In Macaronesia, permits were granted by the Regional Government of the Canary Islands (Exp. 2019/11937; 2020/16669), the Regional Government of the Azores (No. 5/2021/DRAAC), and the Regional Government of Madeira (No. 04/IFCN/2020). In Morocco, sampling was authorised by the *Haut Commissariat aux Eaux et Forêts et à la Lutte Contre la Désertification* (Permit No. 206/2011, 13 January 2011). Fieldwork in the Wold, Germany, was permitted through the Ministerium für Energiewende, Landwirtschaft, Umwelt, Natur und Digitalisierung of Schleswig-Holstein (permit no V 242 - 1765312021 (29-4121)).

## Data accessibility

The raw sequencing data of all individuals included in this study will be available on ENA under the accession number PRJEB71790 upon publication. Information about individuals is available through the supplements.

## Author contribution

CL: conceptualization, data curation, formal analysis, investigation, methodology, project administration, visualization, writing—original draft, writing—review and editing; GL: conceptualization, data curation, formal analysis, investigation, methodology, project administration, visualization, writing—original draft, writing—review and editing; JPT: resources, writing—review and editing; JCI: resources, writing—review and editing; ML: conceptualization, funding acquisition, project administration, supervision, writing—original draft, writing—review and editing

