## Supplementary Information for "From migrants to residents: Genomic insights into adaptive strategies in European robins (*Erithacus rubecula*)"

**Table S1: Summary table for sample location, information and variant calling quality.** Sampling sites are named as follows: WO = Wold, AL = Álava, CO = Covadonga, GU = Guadarrama, LL = Llames, MF =Mirador del Fito, PO = Ponferrada, SD = Sierra de la Demanda, CE = Ceuta, MO = Morocco, SO = Sierra Ojén, EH = El Hierro, LG = La Gomera, LP = La Palma, MA = Madeira, SM = São Miguel, TE = Terceira. We classified individuals as either migratory or resident based on two coincident criteria: 1) the shape and length wing. Thus, sedentary birds had rounded and shorter length wings than migratory individuals; 2) populations were classified as sedentary whether juveniles ringed during the breeding season, were recaptured in the same area during Autumn and/or Winter (see Tellería et al., 2001; Pérez-Tris and Tellería, 2002; De La Hera *et al.*, 2014; De la Hera *et al.*, 2017, for further details). Mean depth is the mean depth calculated over all variants called for that individual, missingness is the amount of missing variants compared to the total number of variants called in the sample set.

| Ringnumber | Population | Phenotype | Grouping | Sex | mean Depth | Missingness |
|---|---|---|---|---|---:|---:|
| 287 | WO | Resident | MG | Male | 36 | 0.0008 |
| R41 | WO | Resident | MG | Male | 18.3 | 0.0030 |
| 269 | WO | Resident | MG | Male | 25.3 | 0.0013 |
| R38 | WO | Resident | MG | Female | 21 | 0.0080 |
| 259 | WO | Resident | MG | Male | 22.3 | 0.0016 |
| A86 | WO | Resident | MG | Female | 9.8 | 0.1910 |
| R13 | WO | Resident | MG | Male | 27.9 | 0.0032 |
| R30 | WO | Resident | MG | Male | 24.9 | 0.0030 |
| R11 | WO | Resident | MG | Female | 17.7 | 0.0042 |
| A29 | WO | Resident | MG | Male | 21.4 | 0.0019 |
| E31 | WO | Resident | MG | Female | 25.7 | 0.0020 |
| N02 | WO | Resident | MG | Male | 20.9 | 0.0076 |
| 202 | WO | Resident | MG | Male | 26 | 0.0013 |
| A45 | WO | Resident | MG | Female | 29.8 | 0.0012 |
| R29 | WO | Resident | MG | Male | 19.9 | 0.0038 |
| C43 | WO | Resident | MG | Male | 22.5 | 0.0020 |
| A46 | WO | Resident | MG | Male | 32.7 | 0.0009 |
| C91 | WO | Resident | MG | Male | 15.9 | 0.0041 |
| 284 | WO | Resident | MG | Male | 24.4 | 0.0015 |
| R42 | WO | Migratory | MG | Male | 22.3 | 0.0018 |
| C96 | WO | Migratory | MG | Male | 20.2 | 0.0022 |
| A23 | WO | Migratory | MG | Male | 21.8 | 0.0023 |
| R85 | WO | Migratory | MG | Female | 15.6 | 0.0080 |
| R16 | WO | Migratory | MG | Female | 21.6 | 0.0096 |
| 265 | WO | Migratory | MG | Male | 22.7 | 0.0016 |
| R90 | WO | Migratory | MG | Male | 24.1 | 0.0036 |
| 260 | WO | Migratory | MG | Male | 15 | 0.0054 |
| N790350 | AL | Migratory | MI | Female | 24 | 0.0016 |
| N790366 | AL | Migratory | MI | Male | 20.2 | 0.0019 |
| N790374 | AL | Migratory | MI | Male | 18.8 | 0.0025 |

| | | | | | | |
|---|---|---|---|---|---|---|
| N790379 | AL | Migratory | MI | Male | 24.1 | 0.0014 |
| L316731 | AL | Migratory | MI | Female | 17.9 | 0.0043 |
| L316741 | AL | Migratory | MI | Female | 24.7 | 0.0016 |
| N238958 | AL | Migratory | MI | Male | 22 | 0.0018 |
| N238963 | AL | Migratory | MI | Male | 18.7 | 0.0024 |
| N238976 | AL | Migratory | MI | Female | 19.9 | 0.0029 |
| N238982 | AL | Migratory | MI | Male | 21 | 0.0018 |
| N238989 | AL | Migratory | MI | Female | 18.8 | 0.0034 |
| PN58 | CO | Migratory | MI | Male | 32.2 | 0.0011 |
| PN60 | CO | Resident | MI | Male | 21.7 | 0.0016 |
| L517419 | GU | Resident | MI | Male | 18.9 | 0.0027 |
| N859812 | GU | Resident | MI | Male | 21 | 0.0022 |
| L517445 | GU | Resident | MI | Female | 22.8 | 0.0021 |
| L517449 | GU | Resident | MI | Female | 20.1 | 0.0029 |
| L517450 | GU | Resident | MI | Male | 16.8 | 0.0039 |
| L517451 | GU | Resident | MI | Male | 19.7 | 0.0025 |
| L517455 | GU | Resident | MI | Male | 24.7 | 0.0047 |
| L316757 | GU | Resident | MI | Female | 16.3 | 0.0057 |
| N238924 | GU | Resident | MI | Male | 22.2 | 0.0016 |
| N238927 | GU | Migratory | MI | Female | 25.8 | 0.0015 |
| PN16 | LL | Migratory | MI | Female | 37.3 | 0.0010 |
| PN17 | LL | Migratory | MI | Male | 25 | 0.0013 |
| PN1886 | MF | Resident | MI | Female | 4.9 | 0.3172 |
| PN1887 | MF | Resident | MI | Male | 13.2 | 0.0067 |
| PN1890 | MF | Resident | MI | Female | 35 | 0.0010 |
| N790410 | PO | Resident | MI | Male | 15.1 | 0.0050 |
| N790415 | PO | Resident | MI | Female | 17.3 | 0.0049 |
| N790416 | PO | Resident | MI | Male | 18.9 | 0.0025 |
| N790314 | SD | Resident | MI | Male | 26.9 | 0.0014 |
| N790315 | SD | Resident | MI | Male | 18.7 | 0.0024 |
| N790317 | SD | Resident | MI | Female | 20 | 0.0027 |
| N790318 | SD | Resident | MI | Female | 19.1 | 0.0030 |
| N790323 | SD | Resident | MI | Male | 14.1 | 0.0052 |
| N790324 | SD | Resident | MI | Male | 16.8 | 0.0029 |
| P2001 | CE | Resident | RC | Male | 29.4 | 0.0013 |
| P2004 | CE | Migratory | RC | Female | 15.2 | 0.0078 |
| P2611 | CE | Migratory | RC | Male | 34.4 | 0.0012 |
| Ro_1208 | MO | Migratory | RC | Male | 24.2 | 0.0024 |
| Ro_1220 | MO | Migratory | RC | Male | 18.2 | 0.0034 |
| L679715 | SO | Migratory | RC | Female | 21 | 0.0029 |

| | | | | | | |
|---|---|---|---|---|---|---|
| L381151 | SO | Migratory | RC | Male | 32 | 0.0012 |
| L679724 | SO | Migratory | RC | Male | 25.8 | 0.0015 |
| L265441 | SO | Resident | RC | Male | 19.5 | 0.0024 |
| L679727 | SO | Resident | RC | Female | 25.6 | 0.0016 |
| L679733 | SO | Resident | RC | Male | 29.4 | 0.0012 |
| L316719 | SO | Resident | RC | Male | 21 | 0.0021 |
| L316720 | SO | Resident | RC | Female | 34.9 | 0.0012 |
| L316723 | SO | Resident | RC | Male | 19.4 | 0.0026 |
| L316724 | SO | Resident | RC | Male | 30 | 0.0013 |
| N328758 | SO | Resident | RC | Male | 23.9 | 0.0019 |
| N328777 | SO | Resident | RC | Female | 35.5 | 0.0011 |
| N185607 | SO | Resident | RC | Male | 25.3 | 0.0014 |
| N328798 | SO | Resident | RC | Female | 18 | 0.0041 |
| N328806 | SO | Resident | RC | Male | 25.9 | 0.0015 |
| N328821 | SO | Resident | RC | Male | 20.6 | 0.0027 |
| N328832 | SO | Resident | RC | Male | 24.7 | 0.0016 |
| N238816 | SO | Resident | RC | Female | 31 | 0.0012 |
| N238818 | SO | Resident | RC | Male | 36.6 | 0.0010 |
| L381194 | SO | Resident | RC | Male | 22.8 | 0.0016 |
| N238820 | SO | Resident | RC | Male | 23.3 | 0.0016 |
| N238821 | SO | Resident | RC | Male | 24.4 | 0.0015 |
| N238823 | SO | Resident | RC | Male | 15.1 | 0.0049 |
| P209420 | EH | Resident | RI | Female | 14.2 | 0.0097 |
| P209423 | LG | Resident | RI | Male | 22 | 0.0031 |
| P209425 | LG | Resident | RI | Male | 20 | 0.0037 |
| P209426 | LG | Resident | RI | Female | 17.5 | 0.0077 |
| 1N87662 | LP | Resident | RI | Female | 32.9 | 0.0010 |
| 1N87667 | LP | Resident | RI | Female | 6.4 | 0.1606 |
| N968323 | LP | Migratory | RI | Male | 12.5 | 0.0088 |
| 2N96568 | LP | Migratory | RI | Male | 20.2 | 0.0021 |
| 2N96569 | LP | Migratory | RI | Male | 28.2 | 0.0012 |
| PE6522 | LP | Migratory | RI | Male | 20.5 | 0.0020 |
| PE7328 | LP | Migratory | RI | Male | 22.5 | 0.0017 |
| 2L55145 | LP | Migratory | RI | Female | 18 | 0.0042 |
| 2L55146 | LP | Migratory | RI | Male | 20.7 | 0.0022 |
| A443764 | MA | Migratory | RI | Male | 17.8 | 0.0045 |
| X40718 | MA | Migratory | RI | Male | 17.7 | 0.0050 |
| A443711 | SM | Migratory | RI | Female | 29.6 | 0.0013 |
| A492510 | SM | Migratory | RI | Male | 21.7 | 0.0036 |
| A492512 | SM | Migratory | RI | Female | 19.4 | 0.0037 |

| | | | | | | |
|---|---|---|---|---|---|---|
| A492528 | SM | Migratory | RI | Female | 16.5 | 0.0073 |
| A492531 | SM | Migratory | RI | Male | 18.4 | 0.0032 |
| No_Ring | SM | Migratory | RI | Male | 21.1 | 0.0045 |
| A492534 | SM | Migratory | RI | Male | 16 | 0.0061 |
| A492540 | SM | Migratory | RI | Male | 17.4 | 0.0049 |
| A407402 | SM | Migratory | RI | Female | 16.9 | 0.0056 |
| A270547 | TE | Migratory | RI | Female | 30.6 | 0.0014 |
| A270548 | TE | Migratory | RI | Male | 24.9 | 0.0017 |
| A270571 | TE | Migratory | RI | Female | 23.2 | 0.0023 |
| A270572 | TE | Migratory | RI | Male | 22.7 | 0.0019 |
| A270576 | TE | Migratory | RI | Male | 26.3 | 0.0014 |
| A270587 | TE | Migratory | RI | Male | 34.7 | 0.0011 |
| A270593 | TE | Migratory | RI | Female | 26.4 | 0.0017 |
| A270599 | TE | Migratory | RI | Female | 17.9 | 0.0045 |
| A270607 | TE | Migratory | RI | Male | 28 | 0.0015 |

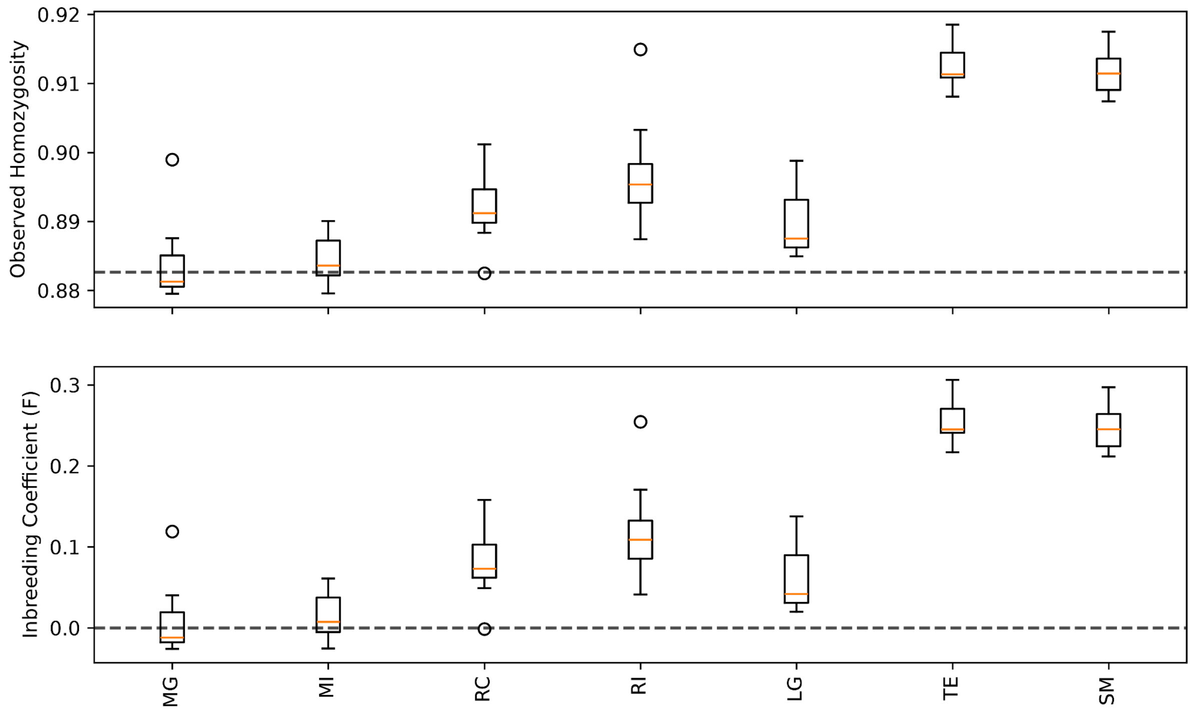


**Figure S1: Expected and observed homozygosity and inbreeding coefficient.** Top: Observed homozygosity of the various populations. Expected homozygosity is shown by the dashed line. In all migratory populations (MG - migratory Germany; MI - migratory Iberia) the observed homozygosity matches expected homozygosity. In resident populations (RC - resident continental; RI - resident Island; LG - La Gomera; TE - Terceira; SM - São Miguel) however, observed homozygosity is much higher than expected. Bottom: this is also reflected in the inbreeding coefficients, which deviate from zero (dashed line) in all resident populations, but is at zero in migrants.

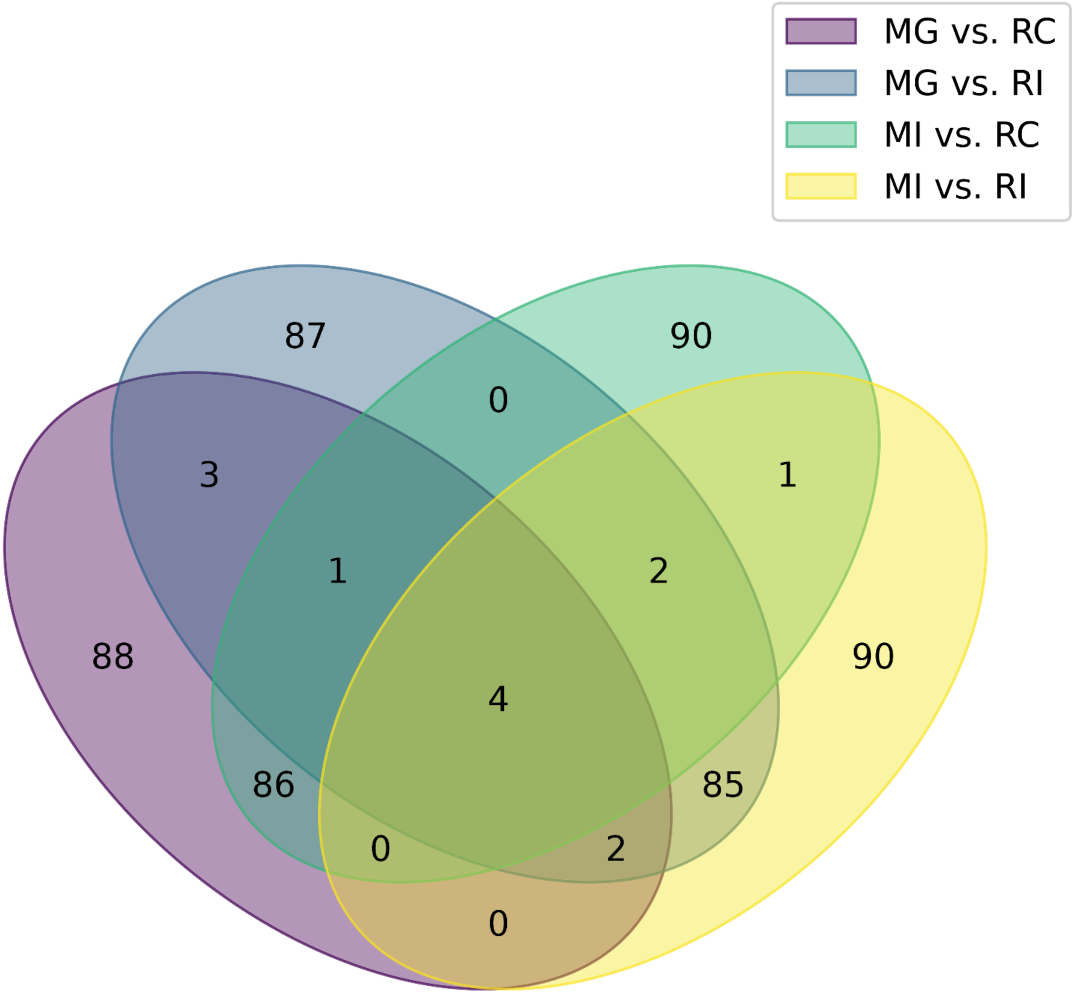


**Figure S2: Venn diagram of outlier windows identified with delta $F_{ST}$ in all migrant-resident comparisons.** Many windows identified are specific to each comparison. Windows identified in two comparisons can be attributed to one population, the one that is present in both comparisons. Here we find that much more identified windows are specific to the resident populations, i.e. are shared between MG (migrant Germany) vs. RC (resident continental), purple and MI (migrant iberia) vs. RC, green. The same is true for RI (resident Iberia) in the overlap between MG vs. RI (blue) and MI vs. RI (yellow).

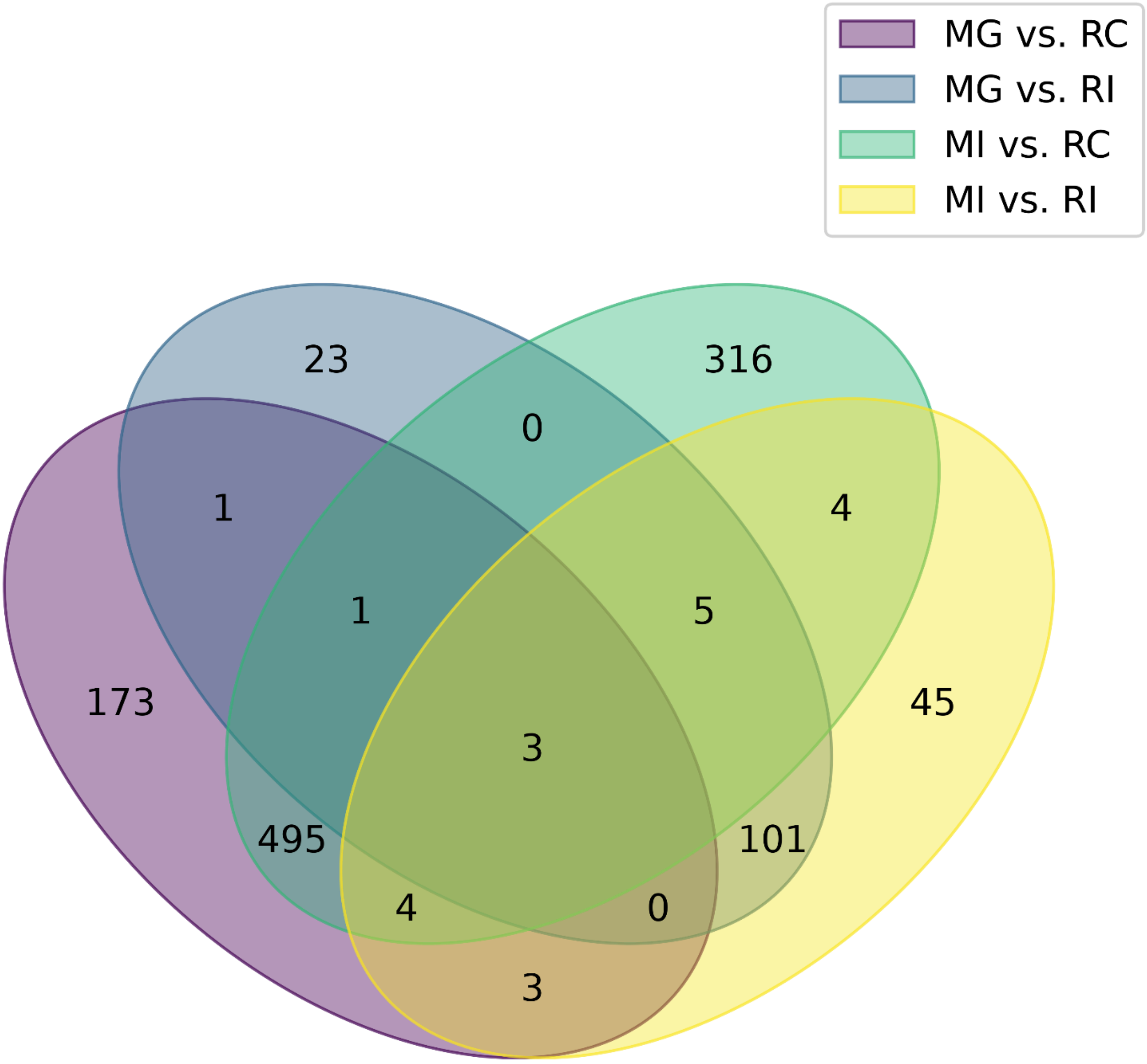


**Figure S3: Venn diagram of outlier windows identified with xp-EHH in all migrant resident comparisons.** This shows the same pattern as figure S2, but additionally shows the difference between resident continental (RC) and resident Island (RI). A lot more windows are specific to RC (see overlap between migrant Germany (MG) vs. RC, purrple and migrant Iberia (MI) vs. RC, green) compared to windows specific to RI (see overlap between MG vs. RI, blue and MI vs. RI, yellow).
